# Single-cell transcriptomic analysis of macaque LGN neurons reveals novel subpopulations

**DOI:** 10.1101/2024.11.14.623611

**Authors:** Shi Hai Sun, Kai Renshaw, John S. Pezaris

## Abstract

Neurons in the lateral geniculate nucleus (LGN) provide a pivotal role in the visual system by modulating and relaying signals from the retina to the visual cortex. Although the primate LGN, with its distinct divisions (magnocellular, M; parvocellular, P; and koniocellular, K), has been extensively characterized, the intrinsic heterogeneity of LGN neurons has remained unexplained. With the development of high-throughput single-cell transcriptomics, researchers can rapidly isolate and profile large sets of neuronal nuclei, revealing a surprising diversity of genetic expression within the nervous systems, such as two types of K neurons (Bakken *et al*., 2021). Here, we analyzed the transcriptomes of individual cells belonging to macaque LGN using raw data from a public database to explore the heterogeneity of LGN neurons. Using statistical analyses, we found additional subpopulations within the LGN transcriptomic population, whose gene expressions imply functional differences. Our results suggest the existence of a more nuanced complexity in LGN processing beyond the classic view of the three cell types and highlight a need to combine transcriptomic and functional assessments. A complete account of the cell type diversity of the primate LGN is critical to understanding how vision works.

## Introduction

The lateral geniculate nucleus (LGN) of the thalamus serves as the main relay through which visual information travels from the retina to the cortex (Sherman & Guillery, 2006; Solomon & Lennie, 2007). Within LGN, visual input is segregated both spatially – distinct areas of the visual field correspond to distinct areas of LGN (Chen et al., 1999; K. A. Schneider et al., 2004) – and functionally (Denison et al., 2014; K. A. Schneider et al., 2004). A bulk of research has focused on determining the precise spatial, functional, and morphological organization of LGN, as a better understanding of LGN’s organization can provide insight into visual processing and has important clinical relevance.

In primates, LGN is composed of distinct layers of functionally similar cell types: magnocellular (M), parvocellular (P), and koniocellular (K). Electrophysiological and morphological studies in primates have determined the specific characteristics of these layers: M neurons, which are large in size and receive rod photoreceptor-derived information, are well-suited for rapid temporal dynamics and detecting motion; P neurons, receiving cone photoreceptor-derived information, are attuned to chromatic changes, making them optimal for processing color and form; and less well-defined K neurons receive short-wavelength ("blue") photoreceptor-derived information (Hubel & Livingstone, 1990; Maunsell et al., 1999; Reid & Shapley, 2002; Tailby et al., 2008; Wiesel & Hubel, 1966).

As methods for electrophysiological investigation improve, evidence has been accumulating for functional heterogeneity within the three primary LGN cell types. For instance, *on*- and *off*-specific responses from the retina are preserved within distinct populations of M, P, and K cells (Ichinose & Habib, 2022; Rousso et al., 2016). K cells have also been shown to express receptive fields with non-standard features, including orientation sensitivity, contrast suppression, and many others (Cheong et al., 2013; Eiber et al., 2018; Solomon et al., 2010). Lastly, a recent electrophysiological analysis identified visually responsive LGN cells that did not exhibit any M, P, or K functional responses, suggesting a functional complexity beyond the M, P, and K classification (Sun et al., 2024). All this evidence points toward the existence of multiple subpopulations of LGN neurons within M, P, and K cell types.

Most recently, single-cell RNA-sequencing technologies have been developed for transcriptomic profiling, enabling the identification of cellular subclasses of cells by comparing the gene expression patterns across large numbers of individual cells within a tissue sample (Lein et al., 2017; Lowe et al., 2017; Ståhl et al., 2016). While past attempts to understand the organization of LGN have relied primarily on morphology and electrophysiology, we hypothesize transcriptomic analysis of LGN neurons will reveal more details about their organization and functional roles in visual processing. For example, single-cell RNA-seq studies in macaque have identified 13 transcriptomically separable excitatory cell classes in V1 (Wei et al., 2022) and more than 60 distinct cell types in the retina (Peng et al., 2019), while only four excitatory cell types were reported in the LGN (Bakken et al., 2021). In the present study, we explore the transcriptomic diversity of macaque LGN neurons using raw data obtained from a published database (Bakken et al., 2021). Statistical analyses of the LGN transcriptomes revealed seven subpopulations of excitatory LGN neurons: two magnocellular populations, two parvocellular populations, and three koniocellular populations. We show that these subpopulations differentially express gene profiles, which imply specific functional differences between these putative subclasses. Our results suggest that there is further nuance in the functional organization of LGN that may be reflected in its role in visual processing.

## Methods

### RNA data

The RNA-seq data consisting of 2,157 macaque nuclei and accompanying metadata were sourced from the Allen Brain Map (Allen Institute for Brain Science, 2022), available at https://portal.brain-map.org/atlases-and-data/rnaseq/comparative-lgn, and originally used in (Bakken *et al*., 2021). For single-cell/-nucleus processing for RNA sequencing and processing, please see Tasic *et al*. (2018); Bakken *et al*. (2018, 2021); Hodge *et al*. (2019) for the complete details. In summary, nuclei were isolated by Fluorescence-Activated Cell Sorting, enriched for neurons with neuronal markers, processed with SMART-seq v4 (Clontech) and Nextera XT (Illumina), and sequenced on HiSeq 2500 (Illumina). RNA-seq reads were aligned to corresponding genomes using the STAR aligner (Dobin et al., 2013), and samples had a median depth of 1.3 million reads/nucleus.

### Preprocessing

Data preprocessing and analyses were performed in R using Seurat v5 (Butler et al., 2018; Hao et al., 2024; Stuart et al., 2019) as Seurat consistently outperforms other modern algorithms (Fu et al., 2025). We used a customized version of the standardized single-cell transcriptomics analysis workflow (Clarke et al., 2021), as described below.

Using the supplied metadata from the (Allen Institute for Brain Science (2022) and Bakken *et al*. (2021), low-quality cells (cells that did not pass quality control criteria: <100,000 total reads, <1,000 detected genes, < 75% of reads aligned to genome, or CG dinucleotide odds ratio > 0.5; Tasic *et al*., 2018) and genes (e.g., uncharacterized or non-informative genes) were removed from all further analyses. All cells were prelabeled as M, P, K (Kap and Kp), GABA (GABA 1-4), or pulvinar, as defined in the metadata. Since this study is a comparison of LGN M, P, and K excitatory cells, GABA and pulvinar cells were removed from further analyses (61% remaining, *n* = 1,309 / 2,157).

### Clustering

M, P, and K cell sets were processed separately using cluster analysis to explore the heterogeneity of LGN neurons. Before clustering, the data was preprocessed and normalized using the Seurat function *SCTransform*, which uses a regularized negative binomial regression model for normalization and analytic Pearson residuals to find top variable genes for downstream analysis (Choudhary & Satija, 2022; Lause et al., 2021). For batch correction, we used *IntegrateLayers* with *HarmonyIntegration*, which integrates cells from multiple datasets (Korsunsky *et al*., 2019). The processed data was then dimensionally reduced using principal component analysis (PCA, via *RunPCA*). Clustering was performed using *FindNeighbors,* K-nearest neighbor (KNN), and *FindClusters*, which uses the Louvain algorithm for cluster identification. The clustering results were visualized using uniform manifold approximation and projection (UMAP, via *RunUMAP*; McInnes *et al*., 2020), and quantitatively evaluated using silhouette scores to assess cluster separation.

To find the optimum parameters to use for the analysis, a global parameter optimization strategy using silhouette scores as the objective metric was used. Specifically, for each of the M, P, and K cell subsets, the optimum number of principal components (PCs), number of features (highly variable genes in *SCTransform*), and clustering resolution (*FindClusters*) were determined.

First, to obtain the optimal number of PCs, the standard deviation of each PC was plotted across a range of feature counts (1,000 to 4,000 in increments of 500) using Seurat’s *ElbowPlot* function. The point where the curve plateaued was selected qualitatively as the optimal number of PCs for each of the M, P, and K cell types. Because the location of the elbow is subjective, we followed the recommendation of the Seurat developers to favor the higher side when choosing this parameter.

Next, a global parameter search was conducted to identify the combination of feature number and clustering resolution that maximized the average silhouette width. The number of features ranged from 1,000 to 4,000 in steps of 100 and the clustering resolution values spanned from 0.1 to 1.0 in increments of 0.02. For each combination, clustering was performed following dimensionality reduction by PCA, using the optimum number of PCs as defined above, and silhouette scores were computed. A Generalized Additive Model (GAM) was then fitted to smooth the silhouette score surface and remove outliers. The parameters corresponding to the peak predicted silhouette score were selected for each cell type group. The final clustering parameters for M, P, and K cell types are specified in the Results section. See Figure 1B for a visualized map of the described clustering workflow.

**Figure 1.**
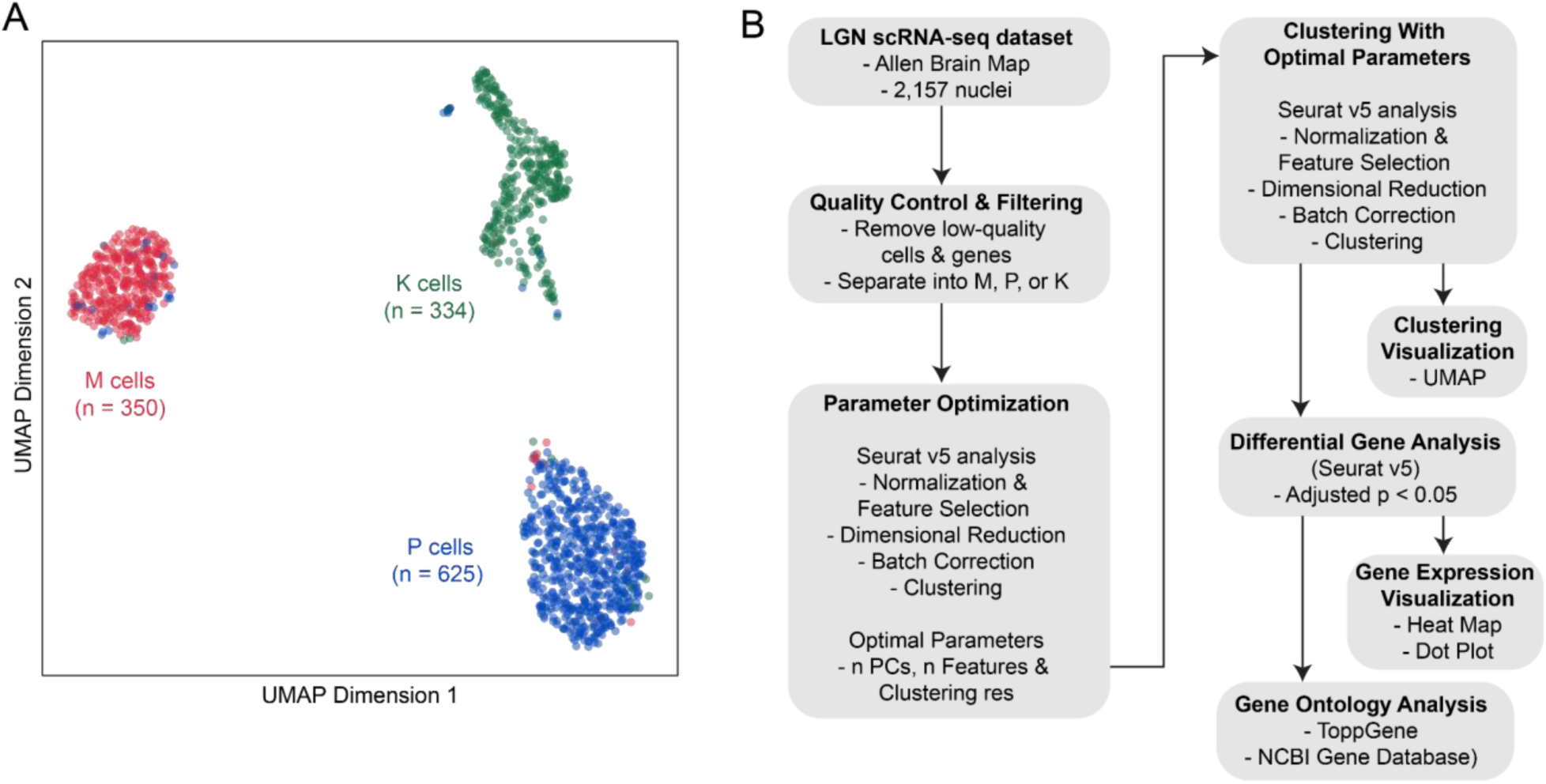
Analysis pipeline of LGN scRNA-seq data, sourced from the Allen Brain Map. **(A)** Scatter plot visualization for each cell (n = 1309) in UMAP space, where the horizontal and vertical axes represent the dimensionality-reduction UMAP coefficients. The magnocellular (M), parvocellular (P), and koniocellular (K) cells are illustrated as red, blue, and green, respectively. **(B)** Flow chart of the clustering and gene analysis. As labeled in the supplied metadata, sets of the M, P, and K cells were individually preprocessed. Each group of cells went through parameter optimization and then clustered using Seurat v5. Differential genes were collected and annotated with each cell population.

### Clustering validation testing

To assess robustness, a sensitivity analysis was performed. This included random downsampling of each dataset to 70-90% of the original cells, as well as ±20% variation in the number of highly variable genes and ±1 variation in the number of dimensions used for clustering. This process was bootstrapped and repeated 1,000 times, and the findings are referenced in the Results and Discussion sections.

To assess the clustering stability we added Gaussian noise directly to the Harmony embeddings, scaled to 20% of the standard deviation (SD) of each embedding dimension. For each cluster group, we performed 1000 iterations of added noise while preserving the original cluster labels, and computed the silhouette score for each perturbed embedding. An empirical *p*-value was calculated as the proportion of noise-affected silhouette scores that were equal to or greater than the silhouette score from the original data. Cluster degradation was defined as the percentage decrease in silhouette relative to the original value.

As an external validation of the clustering approach, an in-silico benchmark was conducted using a publicly available, cell-sorted peripheral blood mononuclear cell (PBMC) single-cell RNA-seq dataset with established ground-truth labels from Wang *et al*. (2019), downsampled from Zheng *et al*. (2017). The dataset was first clustered using the standard Seurat workflow, consistent with methods used in Bakken *et al*. (2021), to broadly identify the immune cell populations. Subsequently, the heterogeneous cluster from the initial analysis was re-clustered using the optimized silhouette-based global parameter search pipeline, as described above. To evaluate stability, the analysis was also repeated under downsampling conditions and across multiple bootstrap iterations. The details and results of this validation experiment are presented in Supplementary Figure 1 and referenced in the Results.

### Differentially expressed genes

Each cluster was profiled through differential gene expression for comparative analysis. Differentially expressed genes/markers (we will use *genes* and *markers* interchangeably) were determined with the Seurat function *FindAllMarkers*, defined with an adjusted *p*-value < 0.05 (Wilcoxon Rank Sum test). We also defined these genes as biologically significant if the log_2_ fold change is greater than 1 (i.e., log2FC > 1 and adjusted *p*-value < 0.05). Gene expression profiles were visualized using heat maps (*DoHeatMap*) and dot plots (*DotPlot*). For the gene ontology analysis, ToppGene (https://toppgene.cchmc.org/enrichment.jsp) and the National Center for Biotechnology Information (NCBI) Gene database (https://www.ncbi.nlm.nih.gov/gene/) were used. As the interpretation of gene ontology results is necessarily speculative, most of this analysis is presented in the Discussion section.

## Results

We analyzed the transcriptomes of 1,309 nuclei from microdissected, anatomically defined regions of the macaque lateral geniculate nucleus (LGN) using RNA-seq data sources from the Allen Brain Map (Allen Institute for Brain Science, 2022; Bakken et al., 2021). The RNA-seq data was preprocessed by removing low-quality cells and unnecessary genes (see Methods), as per the metadata provided by the Allen Institute (Allen Institute for Brain Science, 2022). Like in Bakken *et al*. (2021), the gene expression was quantified as the sum of intron and exon reads, normalized as counts per million (CPM), and log_2_-transformed. Each cell was categorized into one of the predefined clusters (M, P, K) based on the metadata associated with the published data set (Bakken et al., 2021). These cells in their predefined labels are shown in Figure 1A, illustrating the cell-to-cell genetic diversity in a dimensionally reduced UMAP (Uniform Manifold Approximation and Projection) plot.

Given the wide diversity within primate retinal ganglion cells (Dacey, 2004; Peng et al., 2019), our increasing understanding of the complexity of the LGN (Sun et al., 2024) and the visual cortex (Vries et al., 2020; Wei et al., 2022), we believe cellular subtypes within M, P, and K are to be discovered and expect they can be distinguished using transcriptomics. Therefore, each of the M (*n* = 350), P (*n* = 625), and K cell (*n* = 334) groups from the publicly sourced dataset were reanalyzed using an adjusted Seurat analysis pipeline (Figure 1B). Notably, because there are no universally accepted standards for the function parameter selection in the Seurat analysis pipeline (Arbatsky et al., 2025) and the default parameters are inherently arbitrary (I. Schneider et al., 2021), we employed a global optimization approach to optimize for cluster separation quality (Silhouette score) to determine the optimum number of principal components (PCs), number of features, and clustering resolution for each cell type (see Methods).

To confirm the robustness of the clustering pipeline, we first validated the workflow using a ground-truth-labeled PBMC single-cell RNA-seq dataset. The clustering achieved a 90.7% accuracy match to the known labels, with a moderate silhouette score of 0.276 (Supplementary Figure 1). Stability testing through 100 iterations of downsampling and varying the number of features and dimensions yielded a mean cluster identity match of 89.63% (SD = 1.23%), confirming that the pipeline reliably recovers biologically meaningful subpopulations

Our M, P, and K clustering results using the adjusted Seurat analysis pipeline will be described below in threefold: (1) how many within-cluster subpopulations (i.e., clusters within M, P, or K sets) are evident in the data along with the optimized parameters used; (2) what are the significant genetic differences that define these subpopulations; and (3) what functions and interpretations can we infer using genetic annotation from public resources. Much of the interpretation from (3) will be in the Discussion section. Throughout the Results and Discussion, we define significant genes as those with an adjusted *p*-value < 0.05, and biologically significant genes as those with both an adjusted *p*-value < 0.05 and a fold-change > 2 (or log2FC > 1).

However, because the definition of biological significance is somewhat arbitrary (Dalman et al., 2012), and informative genes can exhibit moderate fold changes of less than 2 (St. Laurent et al., 2013), we will consider both significant and biologically significant genes, while explicitly noting fold-change values throughout.

### M cell clustering

Two subpopulations emerged from the set of M cells: M_1_ (*n* = 226) and M_2_ (*n* = 124). The optimized parameters used were 4 PCs, 2800 top features, and a cluster resolution of 0.2 (Figure 2A). After GAM fitting (for smoothing and outlier removal), these parameters produced a silhouette score of 0.231, indicating a low to moderate within-cluster cohesion and inter-cluster separation between M_1_ and M_2_ cells. The two clusters using the optimized parameters can be visualized in the UMAP plot of Figure 2B. Likewise, when only considering the significant genes from these populations (*n* genes = 185), the UMAP plot in Figure 2C illustrates an ideal separation between the two subpopulations with a silhouette score of 0.36, though we acknowledge this is somewhat circular since gene selection was based on differential expression.

**Figure 2.**
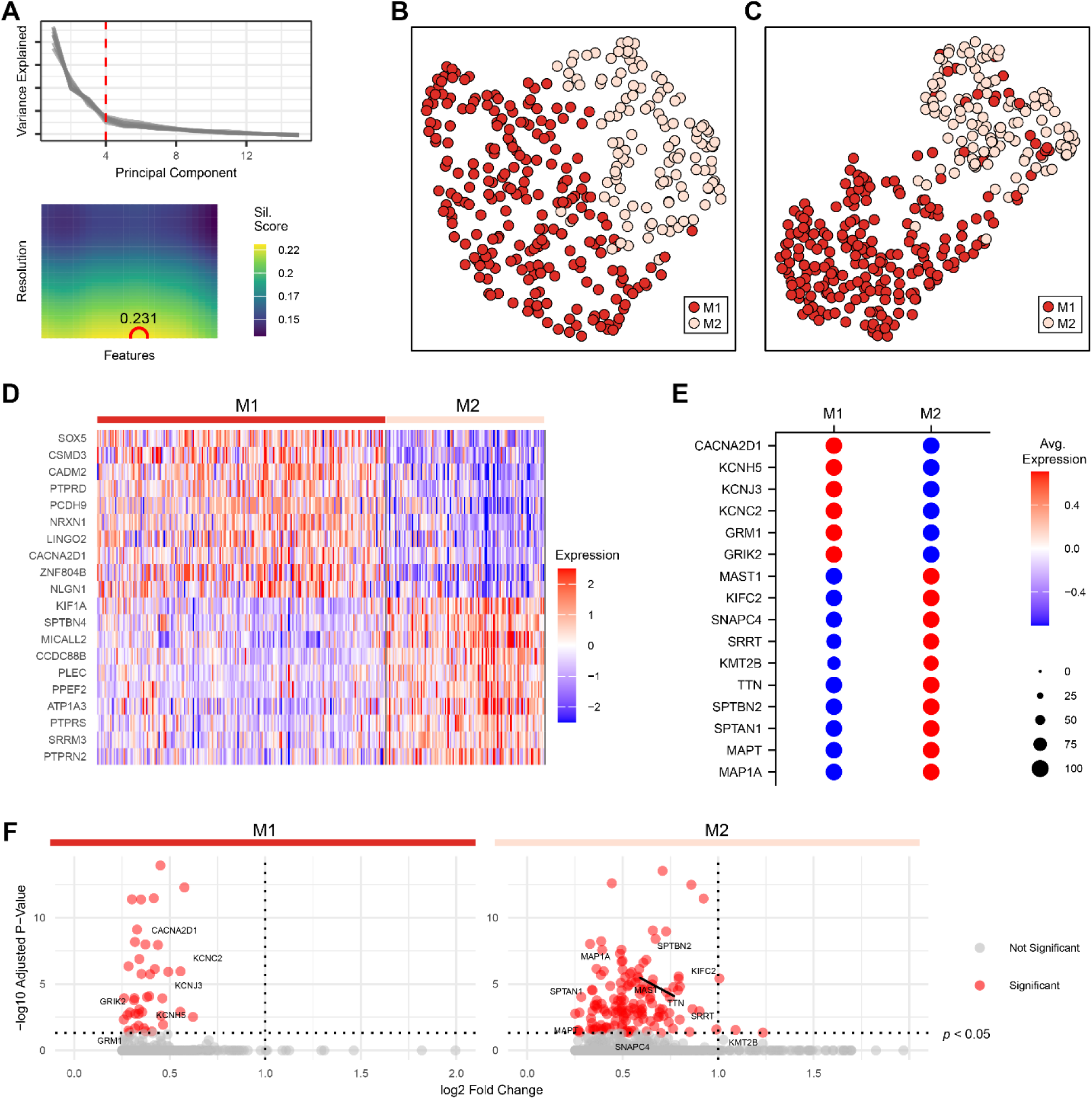
M cell clustering produced two clusters: M_1_ (n = 226, red) and M_2_ (n = 124, light-red). (**A**) Top, elbow plot to qualitatively determine the number of PCs at the elbow (red dashed line). Bottom, heat map of GAM-fitted silhouette scores from varying combinations of number of high variable features and clustering resolution (bottom). The red circle in the heat map indicates the maximum predicted silhouette score. The optimum number of PCs, high variable features, and clustering resolution were used for downstream analysis. **(B)** Scatter plot visualization of M_1_ and M_2_ clusters in UMAP space from the parameters determined in A. **(C)** The same UMAP plot but only using the differentially expressed genes between M_1_ and M_2_ clusters (adjusted *p*-value < 0.05, *n* = 185). (**D**) Heatmap of RNA-seq expression for the top 10 differentially expressed genes (ordered by increasing adjusted *p*-value; positive fold change only) for each cluster. The horizontal and vertical axes represent individual cells and differentially expressed genes, respectively. The colored bars on the top and left of the plot represent the cells and the top 10 differentially expressed for each cluster, respectively. The legend on the right describes the expression levels for each gene and cell. (**E**) Gene expression dot plot showing the expression of notable M_1_ and M_2_ significant markers (adjusted *p*-value < 0.05). Dot diameter indicates the proportion of cells expressing the gene, and color intensity indicates average expression levels. **(F)** Volcano plot of differential gene expression between M_1_ and M_2_ clusters. The x-axis represents the fold change in expression, and the y-axis represents the –log10 adjusted p-value. Genes passing significance threshold (adjusted p-value < 0.05) are represented as red data points. The genes featured in E are labelled for reference.

We also performed stability and sensitivity analyses to assess the robustness of the clustering. Downsampling the number of cells (70-90%), varying the number of highly variable genes (80-120%) and number of PCs (±1), consistently reproduced two M clusters in 71.5% of bootstrap iterations (*n* = 1000), with an average cluster identity match of 86.0 ± 6.0%. This indicates that the optimized parameters are reasonably robust within typical parameter variations. We also tested the clustering parameters under random noise conditions by introducing noise. When adding Gaussian noise to the data (see Methods), no noise-iterated dataset (*n* = 1,000) produced a silhouette score higher than the observed silhouette from the original clustering, resulting in an empirical p-value of 0.008. This indicates that the cluster structure reflects meaningful biological separation rather than random fluctuations. Notably, the silhouette score exhibited only 6% degradation under this noise level, indicating cluster stability.

A total of 185 genes significantly differentiated between M_1_ and M_2_ cells (obtained from Wilcoxon Rank Sum test; see Methods). The genes that contributed the most to the genetic variance between M_1_ and M_2_ cells are illustrated in the heatmap of Figure 2D, showing the expression levels of the most differential genes (top 10 genes from each population ranked by ascending adjusted *p*-value) for each cell. Genes of particular interest from our ontology analysis are shown in the dot and volcano plots (Figure 2E and 2F, respectively).

Generally, from our ontology, M_1_ cells show upregulation of genes associated with action potentials and signal transmission whereas M_2_ cells express cytoskeletal/intracellular transport proteins and genes involved in transcription and post-transcription regulation. Notably enriched genes in M_1_ cells include GRIK2 (ionotropic kainate-type glutamate receptor subunit) and GRM1 (metabotropic glutamate receptor), alongside a diverse array of voltage-gated potassium channel subunits (KCNC2, KCND2, KCNJ3, KCNH5) and the voltage-gated calcium channel auxiliary subunit CACNA2D1. Conversely, M_2_ cells express microtubule-associated proteins such as MAP1A and MAPT, which stabilize dendritic and axonal microtubules respectively, alongside cytoskeletal components SPTAN1, SPTBN2, and TTN. In addition, M_2_ cells express transcriptional regulators including KMT2B, a histone methyltransferase that modulates chromatin architecture and gene expression; SRRT, an RNA-binding protein involved in miRNA processing; and SNAPC4, a component of the snRNA-activating complex essential for snRNA transcription and RNA processing. Finally, M_2_ cells exhibit high levels of genes involved in intracellular transport and cytoskeletal signaling. For example, KIF1A and KIFC2 encode kinesin motor proteins that mediate anterograde and retrograde vesicular transport along microtubules, while MAST1 encodes a microtubule-associated serine/threonine kinase implicated in regulating synapse-associated cytoskeletal dynamics (Tripathy et al., 2018).

Overall, the mild degree of genetic differentiation is consistent with the modest silhouette scores and the lack of sharply distinct clustering in UMAP space. This subtle separation could suggest that M_1_ and M_2_ represent a gradient or continuum of transcriptional subtypes rather than discrete populations. It is also noteworthy that the original work of Bakken et al. (2021) did not identify subtypes within magnocellular neurons, suggesting that the M1/M2 split observed in the current study likely arises from the increased sensitivity of our clustering optimization pipeline (more in the Discussion), although only detecting modest transcriptomic divergence. In summary, while transcriptomic evidence supports the existence of two subpopulations within M cells, the limited biological separation indicates that these likely reflect subtle molecular differences rather than robustly distinct cell types.

### P cell clustering

Similarly, two subpopulations were identified within P cells: P_1_ (n = 334) and P_2_ (n = 291). The optimized parameters were 4 PCs, 3200 top features, and a clustering resolution of 0.26 (Figure 3A) with a 0.225 silhouette score for this clustering, indicating a low to moderate separation as visualized in the UMAP plot (Figure 3B). Restricting to the set of significant genes (n = 692) improves the clustering to a moderate separation with a silhouette score of 0.329 (Figure 3C). Cluster sensitivity testing yielded a moderate cluster reproduction with 55% of iterations matching two P clusters with an average cluster identity match of 82.3 ± 9.4%. Clustering stability testing, adding gaussian noise to the data, resulted in a silhouette score degradation by only 5.88% with no noisy data iteration with a higher silhouette score than the original (empirical *p*-value of 0.000), indicating that the clustering reflects meaningful biological signal rather than noise.

**Figure 3.**
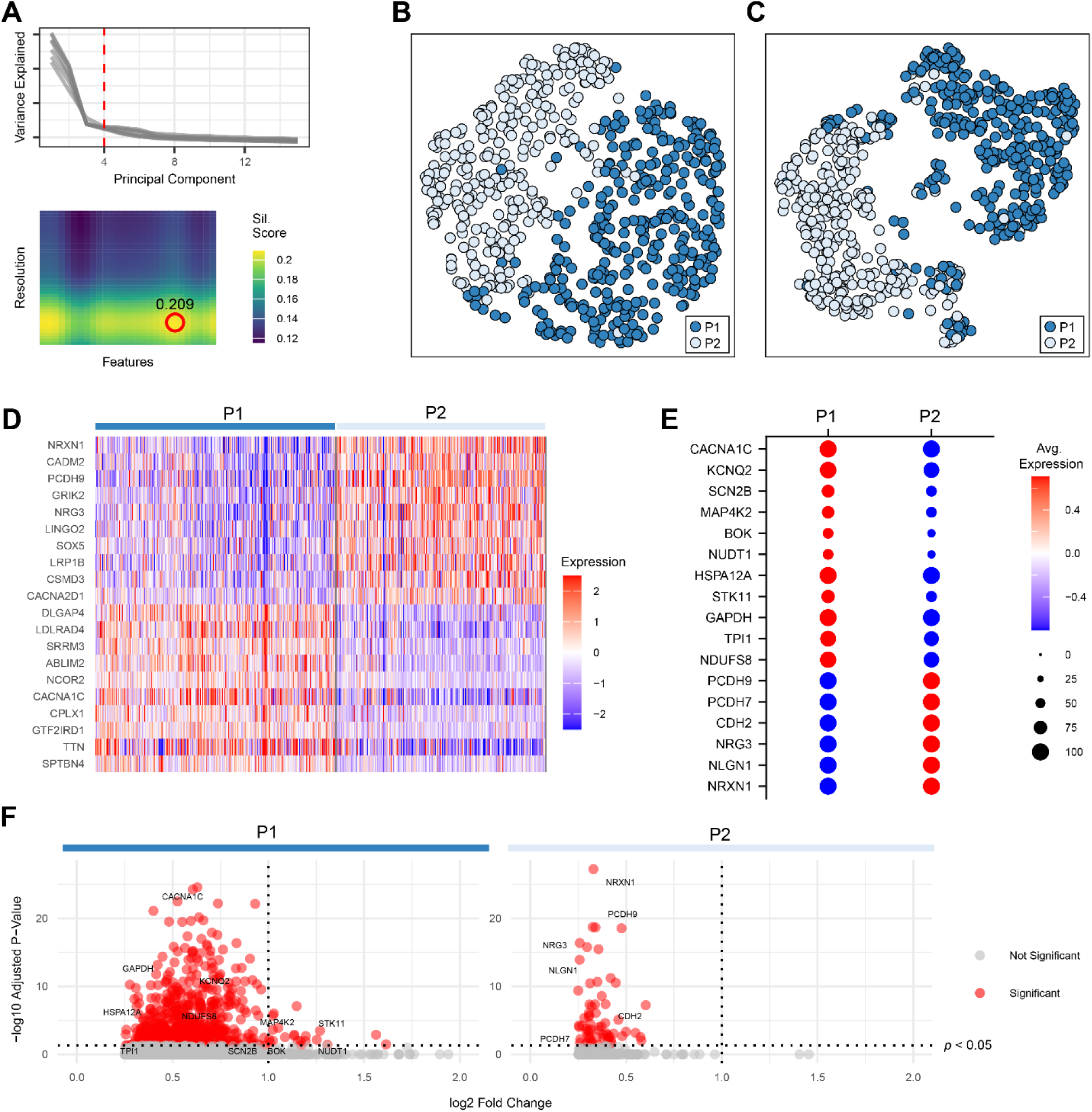
P cell clustering produced two clusters: P_1_ (n = 334; dark blue) and P_2_ (n = 291; light blue). **(A)** Top, elbow plot to qualitatively determine the number of PCs at the elbow (red dashed line). Bottom, heat map of GAM-fitted silhouette scores from varying combinations of number of highly variable features and clustering resolution. The redcircle in the heat map marks the optimal parameters used in downstream analysis. **(B)** Scatter plot visualization of P_1_ and P_2_ clusters in UMAP space using the parameters from A. **(C)** The same UMAP plot restricted to differentially expressed genes between P_1_ and P_2_ (adjusted *p*-value < 0.05, *n* = 692). **(D)** Heatmap of RNA-seq expression for the top 10 differentially expressed genes (ordered by increasing adjusted *p*-value; positive fold change only) for each cluster. Horizontal and vertical axes represent individual cells and genes, respectively. Colored bars denote cluster identity. **(E)** Gene expression dot plot showing expression of notable significant markers in P_1_ and P_2_. Dot diameter represents the proportion of cells expressing the gene; color intensity reflects average expression. **(F)** Volcano plot showing fold change versus –log10 adjusted *p*-value for each gene. Genes meeting both statistical and biological significance are highlighted in red, with markers from **(E)** labeled.

A total of 692 genes were significantly differentially expressed between P_1_ and P_2_ (adjusted *p* < 0.05), with 625 genes enriched in P_1_ and 67 in P_2_. The most significant genetic differences contributing to the P_1_ and P_2_ separation are illustrated in the heatmap of Figure 3D, and genes of particular interest in our ontology are shown in the plots of Figure 3E and 3F.

From our ontology, P_1_ cells express multiple components of the mitochondrial electron transport chain complex I, including NDUFA11, NDUFV1, and NDUFS8, as well as glycolytic enzymes such as TPI1 (triosephosphate isomerase) and GAPDH (glyceraldehyde-3-phosphate dehydrogenase). The mitochondrial GTP-binding protein GTPBP6, implicated in proper mitochondrial translation and oxidative phosphorylation (Lavdovskaia et al., 2020) and STK11, a master regulator of cellular metabolism and energy homeostasis (Huang & Li, 2022), are also highly expressed. In parallel, P_1_ cells express a number of genes that afford protection against oxidative and metabolic stress. These include HSPB1, HSP90AB1, and HSPA12A – heat shock proteins known to stabilize cytoskeletal and synaptic proteins under stress (Dávila et al., 2014; Toth et al., 2010; Wang et al., 2023); NUDT1, which prevents mutagenesis by hydrolyzing oxidized nucleotides; BOK, a BCL-2 family protein involved in mitochondrial dynamics and endoplasmic reticulum stress responses independent of apoptosis (D’Orsi et al., 2016); and MAP4K2, an upstream activator of the JNK pathway, which is a ‘cell death’ pathway involved in cellular stress signaling (Weston & Davis, 2007). Finally, P_1_ cells express a number of ion channels, including SCN2B (voltage-gated sodium channel subunit), KCNT1, KCNAB2, KCNC1, KCNJ12, KCNK3, KCNQ1, KCNQ2 (potassium ion channel subunits), CACNA1G, CACNA1A, CNCBA1B, CACNA1C (voltage-gated calcium channel subunits), and GRIN1 (glutamate NMDA receptor subunit).

By contrast, P_2_ cells express genes such as NRXN1 (neurexin 1) and NLGN1 (neuroligin 1) which, as a pair, function as cell adhesion molecules which bind with each other to establish excitatory/inhibitory synapse specificity (Arias-Aragón et al., 2025; Gomez et al., 2021). NLGN1 specifically promotes excitatory glutamatergic synapse formation (Szíber et al., 2024; Zeidan & Ziv, 2012). NRG3 (neuroregulin) is likewise involved in the formation of excitatory synapses (Müller et al., 2018). CDH2, PCDH7, PCDH9 are all members of the cadherin superfamily, playing roles in axon pathfinding, synaptic targeting, and circuit refinement.

Overall, P_1_ cells show high expression of genes associated with cellular metabolism and energy production, especially aerobic metabolism, as well as a number of genes responsible for protecting the cell from oxidative and metabolic stressors. P_2_ cells, by contrast, express genes important for synaptic regulation and specificity, cell adhesion, and calcium signaling – suggesting an emphasis on connectivity and plasticity.

### K cell clustering

The K cell population exhibited a stronger degree of transcriptional heterogeneity in contrast to P and M cells. Three subpopulations were identified: K_1_ (n = 151), K_2_ (n = 120), and K_3_ (n = 63). The optimized parameters we obtained for K clusters were 4 PCs, 1100 top features, and a clustering resolution of 0.12 (Figure 4A), achieving a stronger silhouette score of 0.401. Using only the significant genes (n = 642) yielded only a slightly higher silhouette score of 0.42 (Figure 4C). The K clustering also produced high stability with an average cluster reproduction match of 92.1 ± 4.5% (*n* = 1000; 14.2% skipped due to cluster mismatch). Noise testing produced an empirical *p*-value of 0.001, with a silhouette degradation of 6.96%, further confirming the robustness of the clustering.

**Figure 4.**
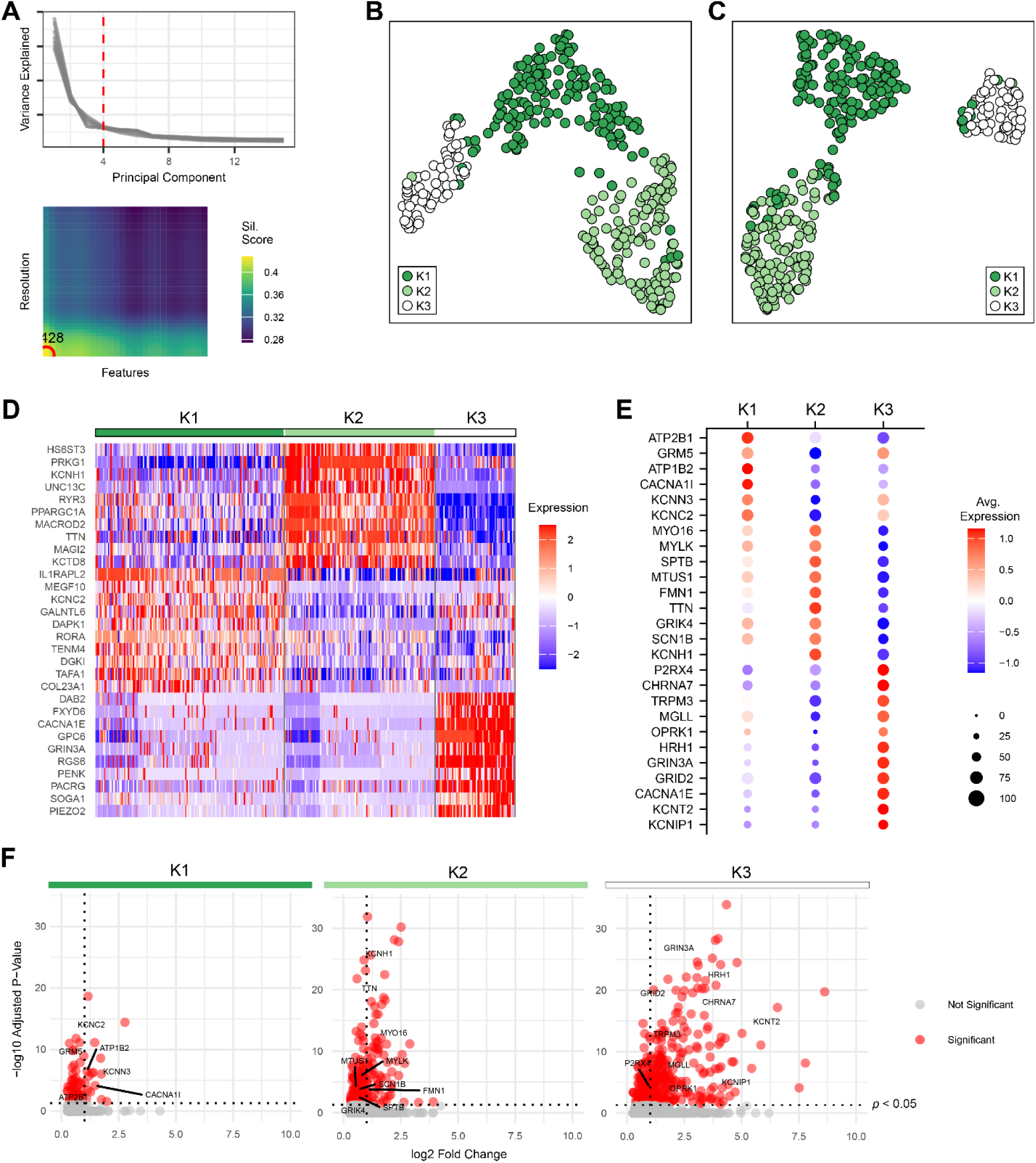
K cell clustering produced four clusters: K_1_ (*n* = 151; dark green), K_2_ (*n* = 120; light green), and K_3_ (*n* = 63; white). **(A)** Top, elbow plot used to determine the number of PCs at the elbow (red dashed line). Bottom, heat map of GAM-fitted silhouette scores across combinations of feature number and clustering resolution. The red circle indicates the optimum parameter set. **(B)** UMAP visualization of K_1_, K_2_, and K_3_ clusters using the parameters in A. **(C)** UMAP using only differentially expressed genes between clusters (adjusted *p*-value < 0.05, *n* = 642). **(D)** Heatmap of RNA-seq expression for the top 10 differentially expressed genes (ordered by increasing adjusted *p*-value; positive fold change only) per cluster. Axes indicate individual cells and gene markers; color bars denote cluster identity. **(E)** Gene expression dot plot showing notable markers in each K cluster. Dot size corresponds to expression frequency; color intensity to average expression level. **(F)** Volcano plot of differential gene expression across clusters. X-axis denotes fold change; Y-axis, –log10 adjusted *p*-value. Genes from **(E)** are labeled.

A total of 642 genes were significantly differentially expressed across the K clusters (adjusted p < 0.05), with 66 enriched in K_1_, 186 in K2, and 390 in K3. Of these, 361 genes met biological significance criteria, including 16 in K_1_, 76 in K2, and 269 in K3 (Figures 4D). This distribution highlights the substantial heterogeneity within the K population, particularly the strong separation of K_3_. Genes of particular interest in our ontology are shown in the plots of Figure 4E and 4F, and described below.

All three K cell subpopulations (K_1_, K_2_, K_3_) express ion channels and glutamate receptors needed for signal transmission and action potential generation, though with subtle functional differences. In addition, coexpression of other genes (such as Ca^2+^-ATPases in K_1_, cytoskeletal binding elements in K_2_, and non-glutamate receptors in K_3_) highlights distinct roles for each subpopulation.

Key genes expressed by K_1_ cells include ion channel subunits such as KCNC2, KCNN3, KCNB2, and CACNA1I, as well as ATP1B2, a subunit of the Na^+^/K^+^-ATPase critical for maintaining resting membrane potential. Notably, K_1_ cells selectively express metabotropic glutamate receptor subunits GRM5 and GRM7, G protein-coupled receptors that mediate slower, modulatory synaptic responses via second messenger cascades. K_1_ cells also express ATP2A3 and ATP2B1, which encode plasma membrane and endoplasmic reticulum Ca^2+^-ATPases involved in calcium extrusion and intracellular calcium homeostasis.

K_2_ cells also express a number of proteins important in signal transmission, including KCNH1, KCNJ3, KCNAB3, SCN1B, SCN9A (sodium and potassium ion channel subunits) and GRIK4, GRIA4 (ionotropic kainate and AMPA glutamate receptors). However, in addition to those proteins important for signal transmission are also a substantial number of proteins involved in cytoskeletal binding, including TTN, TNS1, FMN1, MTUS1, SPTB, MYLK, MYO5B, MYO16, MYO19, MYO14.

The K_3_ population of cells expresses a large number of monoatomic ion channels, channels important for electrical signaling in neurons. These include KCNIP1, KCNA3, KCNQ1, KCNQ5, KCNC4, KCNJ6, KCNK2, KCNK3, KCNH3, KCNT2, KCNK10, KCNMA1 (subunits of different potassium channels, including voltage-gated, sodium-activated, two-pore domain, and inwardly rectifying types), CACNA1E, CACNA1B, and CACNG4 (voltage-gated calcium channel subunits). In addition to these ion channels, this cell population expresses a substantial amount of neurotransmitter receptors for both glutamate – the primary excitatory neurotransmitter in LGN – as well as other modulatory neurotransmitters. These include GRID1, GRID2, GRIN3A, GRM2, GRIK5 (glutamate receptors; NMDA, delta, and kainate type), HRH1 (histamine H1 receptor), OPRK1 (opioid receptor kappa), MGLL (endocannabinoid related lipase), TRPM3 (neurosteroid sensitive transient receptor potential channel; Webster et al., 2020), CHRNA7 (nicotinic acetylcholine receptor subunit), and P2RX4 (purinergic receptor).

In the current analysis, the Kp and Kap cells from Bakken et al. (2021) were analyzed and clustered together due to the relatively small number of Kap cells. Despite this, the clustering from the current analysis consistently grouped Kap cells into K_3_, while subdividing the remaining Kp cells into K_1_ and K_2_. A supplementary figure (Supplementary Figure 2) confirms that K_3_ aligns with the Kap identity (90%, n = 66/73), indicating that the Kap population is preserved and identifiable within the revised clustering framework.

Overall, the strong degree of genetic differentiation within the K population is reflected in the ontology, higher silhouette scores, and clear separation in UMAP space. The dominant transcriptomic signature of K_3_, along with the high number of biologically significant genes, suggests that K_1_, K_2_, and K_3_ represent robust and distinct molecular subtypes. This result refines the earlier findings of Bakken et al. (2021), which identified Kp and Kap subtypes, by further subdividing Kp into K_1_ and K_2_ while preserving Kap as K_3_. The transcriptomic evidence supports the existence of three distinct subpopulations within K cells, indicating a greater degree of heterogeneity in the koniocellular pathway compared to the magnocellular and parvocellular pathways.

### Interspecies and donor variability

To assess whether the observed subpopulations were confounded by interspecies differences or individual subject variability, we performed a qualitative evaluation of UMAP plots for the M, P, and K populations colored by individual animal (*n* = 3) and species (*M. fascicularis* and *M. nemestrina*) (Supplementary Figure 3). As shown in these plots, the major subpopulations are preserved across both subject and species. To quantify this observation, we performed Fisher’s exact tests to evaluate species and subject enrichment within each cluster. No species enrichment was observed in any cluster after Bonferroni correction (*p* > 0.05), indicating that interspecies differences did not drive the clustering. Similarly, no donor enrichment was detected in any of the clusters, except for one koniocellular cluster, K3. K3 exhibited significant donor enrichment (*p* = 0.00023; *p_adj* = 0.0033), likely reflecting the relatively small size of the K3 population. Taken together, these results suggest that the M, P, and K subpopulations are not driven by species- or subject-related differences, and reflect underlying biological distinctions.

## Discussion

Our transcriptomic analysis revealed seven subpopulations of LGN neurons: two populations of magnocellular neurons (M_1_ and M_2_), two populations of parvocellular neurons (P_1_ and P_2_), and three populations of koniocellular neurons (K_1_, K_2_, and K_3_). Although the same dataset was used as Bakken et al. (2021), our study employed several different analytical methods. Notably, we utilized analytic Pearson residuals for normalization (*SCTransform*, Lause *et al*., 2021), Harmony integration for batch correction (*HarmonyIntegration*; Korsunsky *et al*., 2019), and a silhouette-based global parameter optimization pipeline, all of which likely enabled the identification of the nuanced subpopulations in the LGN macaque data.

Examining the gene expression of M, P, and K subtypes elucidates some interesting themes (see Table 1 for summary). As for M and P cells, there appears to be two distinct populations of neurons: first, a population of neurons with genes optimizing for high-throughput signal transmission. In the high-throughput populations of M and P cells, distinct gene expression patterns suggest differences in signaling kinetics, matching the electrophysiologic response profiles observed for each population (Kaplan & Shapley, 1982; Lee & Sun, 2009; Maunsell et al., 1999). The second population of M and P cells shows a striking absence of proteins essential for signal transmission, including ion channels, glutamate receptors, and metabolic genes. Instead, they seem to serve a more modulatory role, expressing genes involved in synaptic plasticity, connectivity, and cytoskeletal structure.

**Table 1.** Summary of ontology analysis.

| Subtype | Key Gene Expression | Functional Implications |
| --- | --- | --- |
| **M<sub>1</sub>** | - Ionotropic (GRIK2) & metabotropic (GRM1) glutamate receptors<br>- Voltage-gated potassium channels (KCNC2, KCND2, KCNJ3, KCNH5)<br>- Voltage-gated calcium channel subunit (CACNA2D1) | - Optimized for rapid, high-fidelity signal transmission<br>- Fast, phasic relay neurons<br>- Fast excitability and spike timing regulation |
| **M<sub>2</sub>** | - Cytoskeletal proteins (MAP1A, MAPT, SPTAN1, SPTBN2, TTN)<br>- Transcriptional regulators (KMT2B, SRRT, SNAPC4)<br>- Kinesin motor proteins (KIF1A, KIFC2)<br>- Microtubule-associated kinase (MAST1) | - Structurally robust, adapted for synaptic plasticity<br>- Modulatory role integrating feedforward and feedback inputs<br>- Supports synaptic remodeling and network activity |
| **P<sub>1</sub>** | - Mitochondrial ETC components (NDUFA11, NDUFV1, NDUFS8)<br>- Glycolytic enzymes (TPI1, GAPDH)<br>- Metabolic regulators (GTPBP6, STK11)<br>- Heat shock proteins (HSPB1, HSP90AB1, HSPA12A)<br>- Ion channels (SCN2B, KCNT1, KCNAB2, etc.)<br>- NMDA receptor subunit (GRIN1) | - High metabolic activity and oxidative stress protection<br>- Tonic firing, high-fidelity relay neurons<br>- Sustained firing patterns suitable for detailed vision |
| **P<sub>2</sub>** | - Synaptic adhesion molecules (NRXN1, NLGN1)<br>- Cadherins (CDH2, PCDH7, PCDH9)<br>- Calcium signaling genes (NRG3) | - Emphasis on synaptic specificity, plasticity, and connectivity<br>- Modulatory role in circuit refinement and activity-dependent changes |
| **K<sub>1</sub>** | - Ion channels (KCNC2, KCNN3, KCNB2)<br>- Na<sup>+</sup>/K<sup>+</sup>-ATPase subunit<br>- Metabotropic glutamate receptors (GRM5, GRM7)<br>- Calcium ATPases (ATP2A3, ATP2B1) | - Slower synaptic response kinetics via metabotropic receptors<br>- High-throughput synaptic activity with calcium regulation |
| **K<sub>2</sub>** | - Signal transmission proteins (KCNH1, KCNJ3, SCN1B, GRIK4, GRIA4)<br>- Cytoskeletal proteins (TTN, TNS1, FMN1, MTUS1, SPTB, MYLK, MYO family) | - Structurally robust with fast conduction properties<br>- Possible projections to extrastriate areas for reflexive/feedforward processing |
| **K<sub>3</sub>** | - Numerous potassium channel subunits (KCNIP1, KCNA3, etc.)<br>- Glutamate receptors (GRID1, GRID2, GRIN3A, GRM2, GRIK5)<br>- Non-glutamate receptors (HRH1, OPRK1, MGLL, TRPM3, CHRNA7, P2RX4)<br>- PENK (proenkephalin) | - Highly electrically active, involved in visual modulation by multiple neurotransmitter systems<br>- Respond to arousal, attention, pain, metabolic, and hormonal states |

K cells do not exhibit the clear high-throughput vs. modulatory division seen in M and P cells. This aligns with numerous studies describing K cells as a more heterogeneous, biochemically unique, and functionally distinct part of the visual system (Hendry & Reid, 2000; Xu et al., 2001). All K neuron populations express proteins important for signal transmission, however the specific signal transmission proteins upregulated in each population gives insight into the response kinetics of each population and highlights the electrophysiologically observed heterogeneity in response kinetics of K cells (Eiber et al., 2018; White et al., 2001; Xu et al., 2001).

Of note, these functional differences of M, P, and K cell populations discussed below come from gene ontology analyses based on a selected subset of expressed genes. While these inferences provide valuable insights, the absence of electrophysiological, morphological, and spatial data limits the ability to comprehensively characterize each cell type.

### Magnocellular

The gene expression profile of M_1_ cells suggests that they are specialized for rapid, high-fidelity signal transmission, consistent with a role as fast, excitable relay neurons adapted to process high-throughput glutamatergic input. The co-expression of both ionotropic (GRIK2) and metabotropic (GRM1) glutamate receptors indicates that these cells are highly responsive to glutamatergic input, likely from retinothalamic projections, through both fast inotropic and slower metabotropic modulatory signaling mechanisms. Kainate receptors such as those encoded by GRIK2 activate within <1 ms at high glutamate concentrations, generating rapid excitatory currents, and rapidly deactivate or desensitize as glutamate levels decline or remain elevated, enabling precise temporal transmission of retinal signals (Lerma et al., 1997). In contrast, metabotropic glutamate receptors encoded by GRM1 are G protein-coupled receptors that modulate downstream signaling pathways, potentially allowing for delayed dynamic tuning of neuronal responsiveness over time (Sugiyama et al., 1987).

High expression of voltage-gated potassium and calcium channel subunits further supports a role in feedforward, high-throughput synaptic transmission. The expression of multiple potassium channel subtypes suggests finely tuned regulation of excitability and spike timing (reviewed in Jan & Jan, 2012). For instance, KCNC2 encodes a high-threshold, fast-activating potassium channel that enables brief, high-frequency action potentials (Yan et al., 2005), while KCND2 contributes to A-type currents that regulate subthreshold excitability, repetitive firing frequency, and back-propagation of action potentials (Connor & Stevens, 1971; Shibata et al., 2000; Zhang & McBain, 1995). CACNA2D1, an auxiliary subunit of voltage-gated calcium channels, enhances calcium channel trafficking and function, supporting efficient neurotransmitter release and dendritic integration. Together, these expression patterns reinforce the hypothesis that M_1_ cells function as fast-acting, phasic relay neurons within the LGN, aligning with the classical view of magnocellular pathways as mediators of high-speed visual processing (Kaplan & Shapley, 1982; Maunsell et al., 1999).

M_2_ cells, in contrast to M_1_ cells, appear structurally robust and preferentially adapted for synaptic plasticity rather than high-throughput signal relay. Their gene expression profile suggests a potential role as modulatory hubs with context-dependent outputs, potentially mediated through dynamic regulation of receptor composition, axonal targeting, or synaptic scaffolding. Notably, M_2_ cells show enriched expression of genes involved in cytoskeletal architecture, intracellular transport, and transcriptional/post-transcriptional regulation. These expression patterns suggest ongoing transcriptional remodeling, possibly in response to activity-dependent plasticity or extra-thalamic modulation.

Principally, these findings support that M_2_ cells are structurally robust. The concerted expression of genes related to cytoskeletal organization, vesicular transport, and transcriptional control – particularly in the relative absence of classical excitability markers such as ionotropic receptors and voltage-gated ion channels – points to two possible, non-mutually exclusive hypotheses regarding M_2_ functionality: first, it’s possible that the gene expression profile of M_2_ cells indicates different action potential kinetics from M_1_ cells. While M_1_ cells are highly excitable, phasic, and fast, M_2_ cells may be more structurally robust, but exhibit slower, more sustained kinetics. One source of evidence supporting this hypothesis comes from many laboratories who have demonstrated that, while P cells demonstrate homogeneously sustained kinetics, M cells contains both transient and sustained neurons (Blakemore & Vital-Durand, 1986; Kaplan & Shapley, 1982; Levitt et al., 2001; Xu et al., 2001). As a second hypothesis, it’s possible that these cells have a specialized role in coordinating synaptic plasticity, integrating feedback and feedforward input, and regulating broader network activity across thalamocortical circuits. M_2_ cells may not be built structurally strong because of size, but instead due to a need to coordinate traffic, form many synaptic connections, and modulate activity broadly. Given the co-expression of many transcriptional regulatory proteins, these cells may serve a more integrative role within the LGN, optimized for modulatory functions rather than rapid signal relay.

Overall, M cells appear to have two functionally distinct populations. M_1_ cells are the prototypically-understood definition of magnocellular neurons. They appear to be the work-horses of electrical transmission, but simple relay neurons, transmitting information from retina to cortex with minimal modulation or refinement. Their presumed response kinetics are the classically understood fast response kinetics and high temporal frequencies associated with M cells. M_2_ cells seem to be more modulatory players. With an absence of proteins important for excitability, action potentials, or synaptic transmission, these are structurally robust neurons, possibly with extensive networks of dendrites or axons. Their high expression of proteins involved in transcriptional and post-transcriptional suggests that they have a role in activity- or state-dependent modulation of neural activity. As opposed to simple relay neurons, M_2_ neurons may be responsible for visual processing of retinofugal input to LGN.

### Parvocellular

P_1_ cells exhibit a gene expression profile marked by high metabolic activity and enhanced resilience to oxidative and mitochondrial stress, suggesting specialization for sustained, energy-demanding function. Among the most biologically significant markers are numerous enzymes involved in core metabolic pathways and cellular stress regulation. Notably, P_1_ cells express multiple components of the mitochondrial electron transport chain complex I, as well as glycolytic enzymes. Mitochondrial GTP-binding protein GTPBP6, implicated in proper mitochondrial translation and oxidative phosphorylation (Lavdovskaia et al., 2020) and STK11, a master regulator of cellular metabolism and energy homeostasis (Huang & Li, 2022) support the interpretation that P_1_ neurons have high, sustained energy demands, relying heavily on aerobic respiration to maintain function under prolonged activity. Finally, P_1_ cells express a number of genes, such as heat shock proteins, NUDT1, BOK, and MAP4K2, that confer protection against oxidative and metabolic stress.

To further support the hypothesis that P_1_ cells function as a high-throughput population of relay neurons, these cells exhibit robust expression of genes involved in action potential generation and synaptic transmission, paralleling the molecular profile of M_1_ cells. These include a voltage-gated sodium channel subunit, potassium ion channel subunits, voltage-gated calcium channel subunits, and glutamate NMDA receptor subunit.

This gene expression profile supports the hypothesis that P_1_ cells are metabolically robust relay neurons. Like M_1_ neurons, P_1_ neurons may serve high-throughput relay functions, however the distinctive upregulation of genes associated with oxidative metabolism, mitochondrial integrity, and stress resilience in P_1_ cells suggests a tonic firing profile rather than the fast, phasic signaling characteristic of M_1_ cells. This aligns with classical physiological descriptions of parvocellular neurons as exhibiting sustained firing patterns in support of detailed, high-fidelity, continuous visual processing (Lee & Sun, 2009; Maunsell et al., 1999).

While P_1_ cells express proteins suggestive of high-throughput signal transmission, P_2_ cells demonstrate gene expression important for synapse formation and specificity, cell adhesion, and calcium signaling, with a notable absence of proteins related to signal transmission – such as ion channels or neurotransmitter receptors. These features suggest a highly plastic and well-connected population of neurons which, like M_2_ neurons, may function as a connectional hub or integrator, rather than a high-throughput information relay. Overall, genes such as NRXN1 and NLGN1, as well as multiple members of the cadherin superfamily, suggest that P_2_ cells play a role in precision wiring and synaptic specificity, forming highly targeted connections. The expression of activity-regulated structural proteins (e.g., neurexins and neuroligins) may indicate a role in activity-dependent circuit adaptability (Jüngling et al., 2006).

Like M cells, P cells also have two functionally distinct populations. P_1_ cells (similar to M_1_ cells) are the high-throughput signal transmitters with gene expression suggestive of substantial metabolic needs, as well as protection from metabolic stress. These features suggest cells which fire tonically, specific for high-fidelity, high-detail vision. P_2_ cells, like M_2_ cells, appear to play a more modulatory, information-integration role. They seem to be well connected, with proteins specific for synaptic specificity and involved in activity-dependent structural modulation and regulation.

### Koniocellular

Similar to M_1_ and P_1_ cell populations, K_1_ cells exhibit enriched expression of genes associated with action potential generation and synaptic transmission. However, the specific repertoire of signal transmission proteins in K_1_ cells offers insight into their distinct response kinetics and functional specialization relative to M and P neurons. The absence of ionotropic glutamate receptors and the exclusive expression of metabotropic receptors suggest that K_1_ cells may exhibit slower synaptic response kinetics, characteristic of the longer response latencies associated with K cells (Pietersen et al., 2014; Xu et al., 2001). These studies also noted highly heterogeneous response kinetics associated with K cells, which is demonstrated by the diversity of different potassium, calcium, sodium, and glutamate receptor types expressed in K_1_, K_2_, and K_3_ cells. The expression of ATP2A3 and ATP2B1, calcium clearance pumps, supports the idea that K_1_ cells are equipped to handle high-throughput synaptic activity while minimizing calcium-induced cytotoxicity.

In K_2_ cells, the presence of cytoskeletal binding proteins suggests that these neurons, like M_2_ neurons, are structurally robust. With the coexpression of fast sodium and potassium channel subunits and ionotropic glutamate receptors, one hypothesis is that the upregulation of structural proteins reflects thicker axons with faster conduction velocities, perhaps with projections to extrastriate areas. This implies that K_2_ cells could have a role in reflexive behaviors or feedforward signalling, projecting to areas requiring fast, high-bandwidth processing, and could possibly be the basis for observed projections from K layers to MT (Casagrande, 1994; Sincich et al., 2004).

The K_3_ population of cells expresses many proteins important for electrical signaling in neurons as well as a diverse population of glutamate and non-glutamate neurotransmitter receptors. The increased expression of these genes suggests that this population of cells is highly electrically active and may be the site of visual modulation by non-glutamate neurotransmitter systems within LGN.

Like K_1_ and K_2_ cells, the expression of ion channels and glutamate receptors indicate an active role in signal transmission. The diversity of glutamate receptor types (NMDA, kainate, and delta type ionotropic, as well as metabotropic) and ion channels expressed in K_3_ continue to support the electrophysiological findings that K cells overall demonstrate highly heterogeneous response characteristics. The increased expression of potassium ion channels relative to other K subtypes implies that K_3_ cells may demonstrate high temporal frequencies similar to M cells – a point to which we will return. However, what is unique about K_3_ cells is the vast expression of non-glutamate neurotransmitter receptors. These include receptors specific for histamine, opioids, acetylcholine, extracellular ATP, and lipids/steroids. The expression of these receptors suggest that K_3_ cells are under tight top-down and subcortical control; they respond to arousal, attention, alertness, pain, and inflammation, dynamically shifting their output depending on internal brain states (Javadi et al., 2015; Jin et al., 2002; Noseda et al., 2017; Yang et al., 2015). In addition, these neurons may adapt their function in response to steroid hormones, metabolic shifts, or stress, potentially linking vision or sensory processing with hormonal states.

The final remark regarding K_3_ cells is the expression of PENK (proenkephalin). Bakken et al. (2021) previously performed single-cell RNA sequencing on macaque LGN cells and showed two populations of K cells, Kp and Kap. Kp cells selectively expressed PENK and were enriched in ventral K layers K1 and K2, adjacent to M cells. This association suggests that the K_3_ cells we identified were the same Kp cells identified by Bakken et al., or a subpopulation thereof, and may help to explain the commonality in response characteristics between M and K_3_ cells. In fact, research in the K1 and K2 layers (where these Kp/K_3_ cells are enriched) has shown that these cells demonstrate large receptive fields with sensitivity to contrast and high temporal frequencies – very similar to M cell receptive fields (Solomon et al., 2002; White et al., 2001; Xu et al., 2001).

## Conclusion

Overall, the transcriptomic evidence supports a refined view of LGN organization, extending beyond the classical M, P, and K classification. The presence of modest subtypes in M and P pathways and strong molecular heterogeneity in K cells suggests that LGN neurons may play a more diverse and functionally specialized role in visual processing than traditionally held. This heterogeneity could reflect differential involvement in non-visual modulation, temporal dynamics, or attentional processes.

Future studies that combine transcriptomics with electrophysiological or anatomical data, such as spatial transcriptomics or patch-seq, will be necessary to validate these subpopulations and clarify their roles. Furthermore, matched functional-transcriptomic datasets would be essential to determine whether any of the transcriptomic clusters observed here correspond to the functionally distinct cells described in recent electrophysiology experiments, to draw further conclusions about the physical organization of the LGN and its rich variety of cell types.

## Supplementary Materials

**Supplementary Figure 1.**
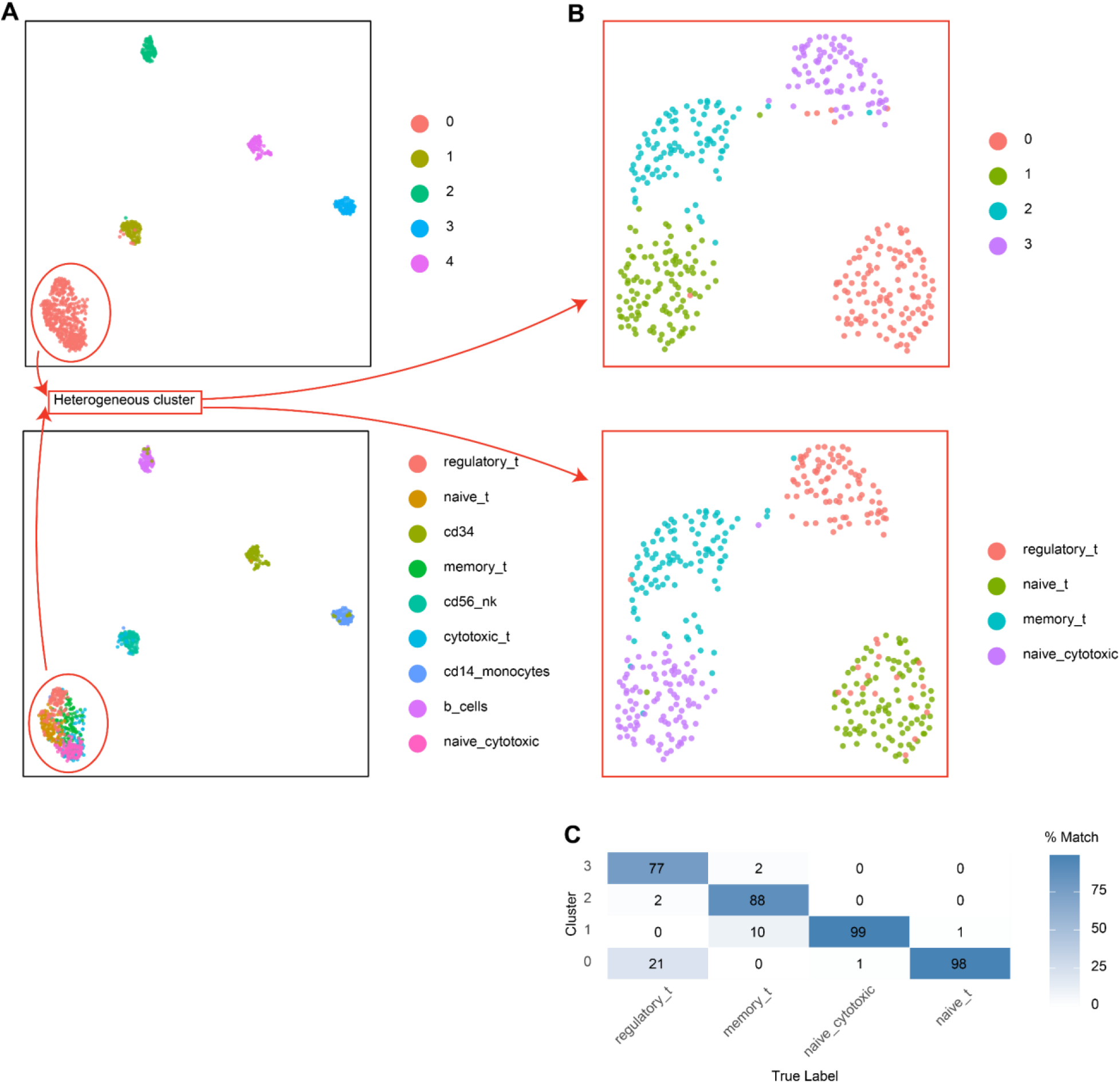
Testing the clustering analysis pipeline on ground truth PBMC data. **(A)** UMAPs of clustering using the default Seurat parameters. This produced 5 clusters, including one heterogeneous cluster (bottom-left), when compared to the true labels (bottom UMAP). **(B)** The heterogeneous cluster in A was reclustered using the same clustering pipeline as described in this study (top UMAP). Bottom, the same UMAP plot but labeled using the growth truth labels. **(C)** Confusion matrix showing match (%) between clusters (y-axis) and true label (x-axis). Overall, the clustering pipeline produced a 90.73% match with the ground truth data.

**Supplementary Figure 2.**
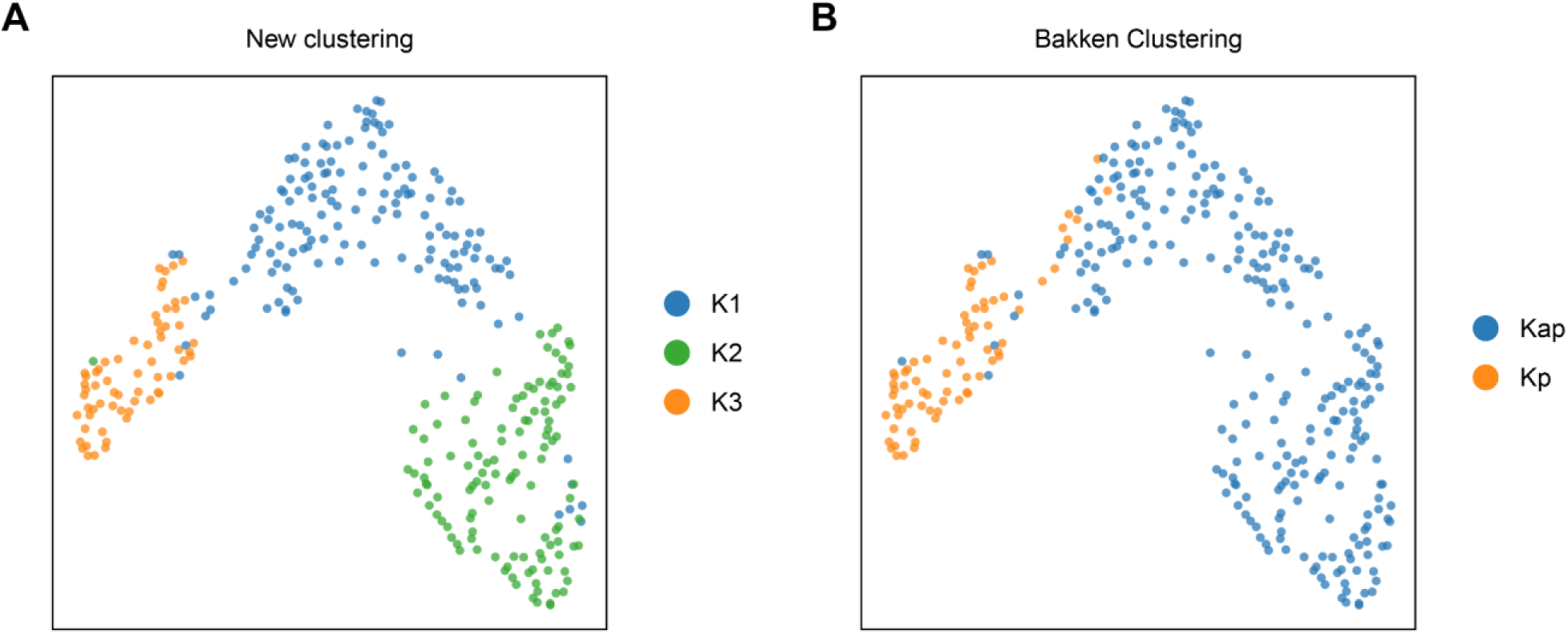
K clustering from the current study **(A)** and from the Bakken et al. (2021) study **(B)**. K3 in the current study is analogous to Kp in the Bakken study with an 86.3% match (*n* = 63/73).

**Supplementary Figure 3.**
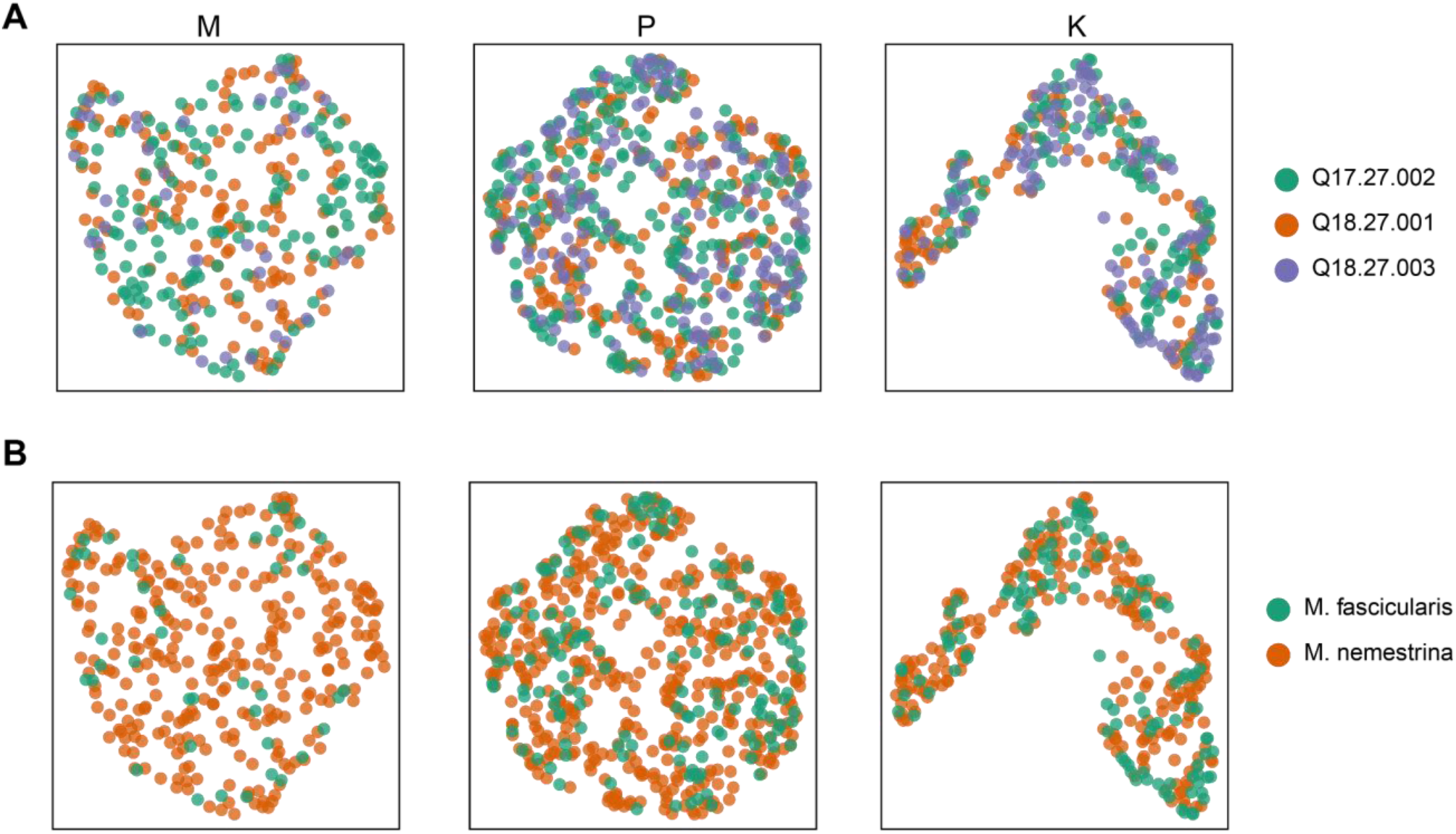
The UMAPs of M (left), P (middle), and K clusters (right) with the data labelled by animal id (**A**, n = 3) and species (**B**, n = 2).

## Data and Code Availability

The R scripts used for data preprocessing, clustering, and analysis in this study are available online at https://github.com/shihaisun-scott/sun_LGN_macaque_transcriptomics.

